# Cellular dynamics of endosperm development in *Arabidopsis thaliana*

**DOI:** 10.1101/2022.04.01.485647

**Authors:** Mohammad Foteh Ali, Ji-Min Shin, Umma Fatema, Daisuke Kurihara, Frédéric Berger, Ling Yuan, Tomokazu Kawashima

**Author notes:** Correspondence to Tomokazu Kawashima. These authors contributed equally: Mohammad Foteh Ali, Ji-Min Shin.

## Abstract

After double fertilization, the endosperm in the seeds of many flowering plants undergoes repeated mitotic nuclear divisions without cytokinesis, resulting in a large coenocytic endosperm that then cellularizes. Growth during the coenocytic phase is strongly associated with the final seed size; however, a detailed description of the cellular dynamics controlling the unique coenocytic development in flowering plants has remained elusive. By integrating confocal microscopy live-cell imaging and genetics, we have characterized the entire development of the coenocytic endosperm of *Arabidopsis thaliana* including nuclear divisions, their timing intervals, nuclear movement, and cytoskeleton dynamics. Around each nucleus, microtubules organize into aster-shaped structures that drive F-actin organization. Microtubules promote nuclear movement after division while F-actin restricts it. F-actin is also involved in controlling the size of both the coenocytic endosperm and mature seed. Characterization the of cytoskeleton dynamics in real-time throughout the entire coenocyte endosperm period provides foundational knowledge of plant coenocytic development, insights into the coordination of F-actin and microtubules in nuclear dynamics, and new opportunities to increase seed size and our food security.

## Introduction

Flowering plants perform a unique double fertilization^1,2^. The pollen tube contains two sperm cells, one of which fertilizes the egg cell and the other fertilizes the central cell to generate the embryo and endosperm in the developing seed, respectively^3–5^. The endosperm serves as a nourishing tissue for the developing embryo during the early phase of seed development. In many monocots such as rice, wheat, and corn, the endosperm persists until maturation and stores carbohydrates and proteins, which are the primary food source for humankind^5–7^, whereas in dicots such as beans and *Arabidopsis thaliana*, the endosperm is consumed by the embryo during subsequent seed development^4^. The endosperm not only holds great agricultural importance but also has an essential role in the evolutionary success of flowering plants.

In *Arabidopsis*, endosperm development follows four developmental phases: coenocyte, cellularization, differentiation, and cell death^3,8^. The coenocyte development starts immediately after fertilization of the central cell, which undergoes several rounds of nuclear division without cytokinesis^9^. The endosperm enlarges and differentiates into the micropylar endosperm (MCE), the chalazal endosperm (CZE), and the peripheral endosperm (PEN). After rounds of repeated mitotic nuclear divisions, the coenocytic endosperm starts to cellularize from the MCE toward PEN and it remains uncellularized in the CZE^9^. The timing of the transition from the coenocytic endosperm to cellularized endosperm determines the final seed size; shorter coenocytic endosperm periods or precocious endosperm cellularization results in relatively smaller seeds, whereas longer coenocytic periods or delayed endosperm cellularization are associated with enlarged seeds^10–17^. It remains largely unknown what cellular dynamics occur in the early phase of endosperm growth and how they control this unique coenocytic development.

In both plants and animals, cytoskeletal structures such as actin filaments (F-actin) and microtubules (MTs) regulate many fundamental cellular processes^18–25^, including those in plant reproduction such as pollen tube growth and guidance, sperm nuclear migration, and asymmetric division of the zygote^18–20,26–28^. Immunostaining studies of coenocytic endosperm at interphase revealed a nucleus-based radial MT (aster-shaped) system that organizes cytoplasm into nuclear-cytoplasmic domains^21–24^. F-actin shows reticulate patterns during the mitotic phase, but its function has not been reported^21,23^. The advancement of live-cell imaging using confocal microscopy has enabled us to visualize F-actin and MT dynamics with the nuclei in real time. We performed both pharmacological and genetic analyses in *Arabidopsis* to characterize the complete development of coenocytic endosperm, including the details of nuclear movements, nuclear divisions, and division timings from fertilization until endosperm cellularization. Immediately following the nuclear divisions, aster-shaped MTs around each nucleus become a foundation for F-actin aster organization and both MTs and F-actin are indispensable for nuclear organization during the coenocytic phase of endosperm development. Our results also showed that the manipulation of F-actin dynamics in the coenocytic endosperm affects the final seed size without altering the timing of endosperm cellularization, revealing a new regulatory mechanism for controlling seed size.

## Results

### Live-cell imaging reveals coenocytic endosperm nuclei dynamics

We performed time-lapse confocal microscopy to monitor nuclear movements and divisions during the coenocytic phase in *Arabidopsis* endosperm *(proFWA::H2B-mRuby2*^18^). The division of the primary endosperm nucleus was observed approximately 2-3 hours from the observation start (Fig. 1a, g and Extended Data Table 1 and Video 1), consistent with previous reports^9,29,30^. The first four nuclear divisions, occurring between 0 and 1 DAP (days after pollination), were synchronous and rapid with 3 to 6 hour intervals (Fig. 1a-d, g and Extended Data Table 1 and Video 1). As the micropylar-chalazal axis extended and bent at the chalazal end, nuclei moved toward both micropylar and chalazal poles and were positioned in an equidistant manner (Fig. 1b-c). After the 3^rd^ division (8 nuclei), two nuclei at the chalazal side moved further into the chalazal pole (Fig. 1c), founding the CZE^9^.

**Figure 1:**
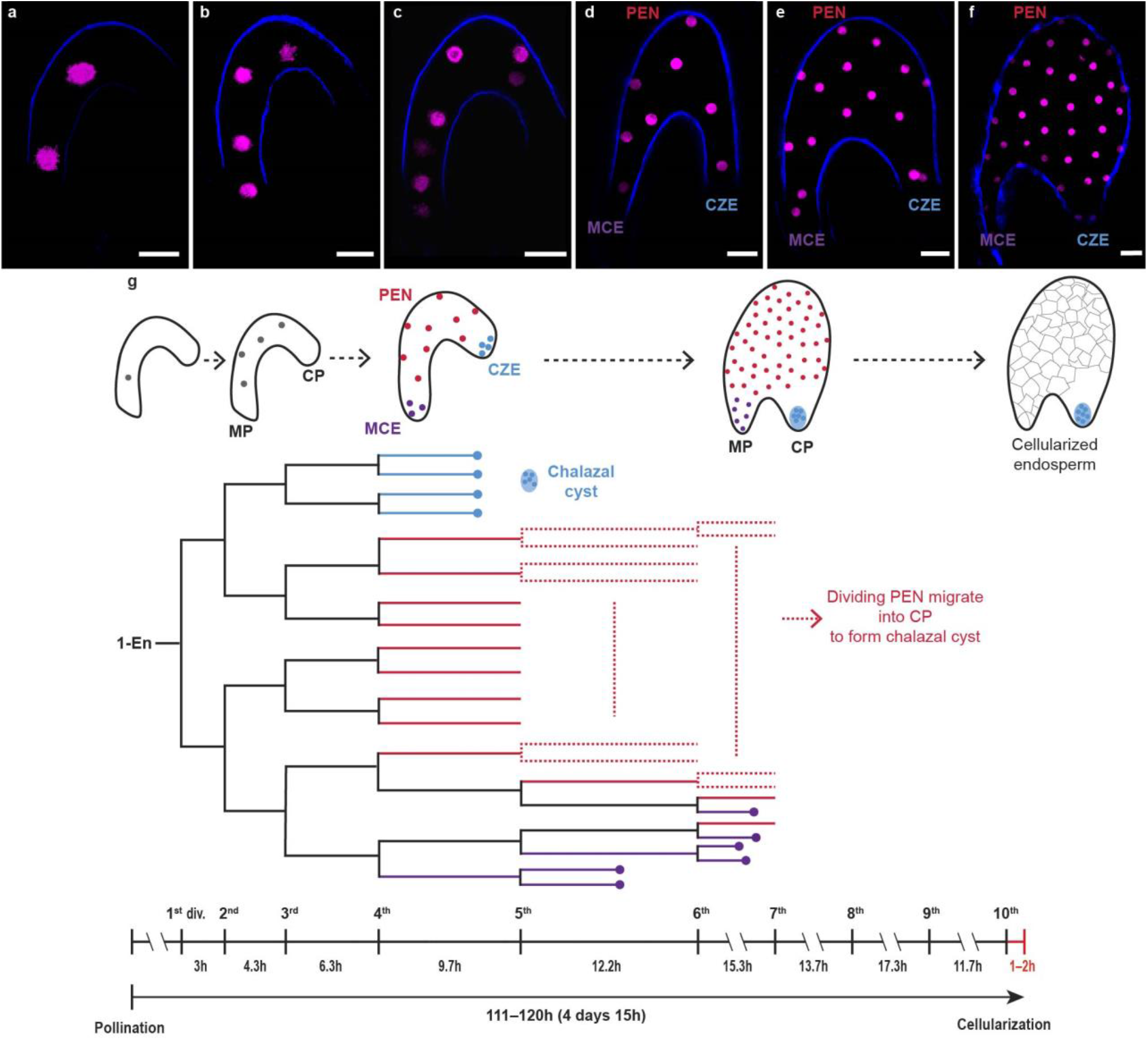
Dynamics of coenocytic endosperm development. **a-f**, Z-projected confocal images of coenocytic endosperm nuclei (magenta, *proFWA::H2B:mRuby2*) after the 1^st^ to 5^th^ (**a-e**), and 7^th^ or 8^th^ divisions (**f**). Autofluorescence (blue) marks the coenocytic endosperm border. Scale bar, 20 µm. **g**, Schematic representation of coenocytic endosperm development (top), the lineage of endosperm nuclei fate from primary endosperm to endosperm cellularization (middle), and nuclear division intervals (bottom). The dots in the lineage indicate nuclei which did not divide further. The numbers of seeds observed for 1^st^ division to 10^th^ division intervals are shown in Extended Data Figure 3. MP, micropylar pole; CP, chalazal pole; CZE, chalazal endosperm; MCE, micropylar endosperm; PEN, peripheral endosperm. **a-g** correspond to Video 1.

After the 4^th^ division (16 nuclei), the three endosperm sub-regions, the MCE, the PEN, and the CZE, followed distinct nuclear division patterns (Fig. 1d, g and Video 1). The enlargement of the PEN began and all PEN nuclei maintained active synchronous division until cellularization (Video 1). After the 5^th^ division, 3-4 nuclei were moved to the end of the MCE, and 1-2 nuclei among these MCE nuclei ceased dividing (Fig. 1e, g and Video 1). In the CZE, the two nuclei that moved to the chalazal pole after the 3^rd^ division divided once more, and these four became the foundation of the multinucleate chalazal cyst (Fig. 1d,e, g and Video 1). The cyst enlarged during development and continuously incorporated nuclei from the PEN (Fig. 1g and Video 1).

Nuclear division intervals in the PEN and the MCE regions after the 4^th^ division became successively longer (1 DAP) with the progression of divisions; the 3^rd^, 4^th^, and 5^th^ divisions took 4-5, 6-7, and 8-9 hours, respectively (Fig. 1g). From the 6^th^ division, the endosperm nuclei division intervals were 14-18 hours until the 9^th^ division (5 DAP), and the last division took 11-12 hours (Fig. 1g). In total, ten nuclear divisions in the PEN were observed before cellularization in our live-cell imaging system (Fig. 1g and Extended Data Table 1 and Video 1). Cellularization was initiated 1-2 hours after the 10^th^ division (5 DAP; Fig. 1g and Video 1), starting from the MCE to PEN. The CZE remained uncellularized, consisting of chalazal nodules and cyst. Both *in planta* (Extended Data Fig. 1a) and semi-*in vivo* (Fig. 1g), the total duration of all nuclear divisions in coenocytic endosperm development was approximately 5 DAP, demonstrating that our live-cell imaging system reflects the development of young *Arabidopsis* seeds *in planta*.

### F-actin generates unique aster-shaped structures around each nucleus of the MCE and the PEN and controls the nuclear position

To understand how the cytoskeleton is involved in the dynamics of coenocytic endosperm nuclei, we monitored coenocytic endosperm F-actin dynamics (*proFWA::Lifeact-Venus*^19^; Fig. 2). There is a constant inward movement of F-actin meshwork for sperm nuclear migration in the central cell upon fertilization (Fig. 2a) and this inward movement disappears after successful fertilization^19^. Until after the 2^nd^ endosperm nuclear divisions, F-actin retained a reticulate cable network throughout the cell that enmeshed each endosperm nucleus (Fig. 2b, c and Extended Data Fig. 2).

**Figure 2:**
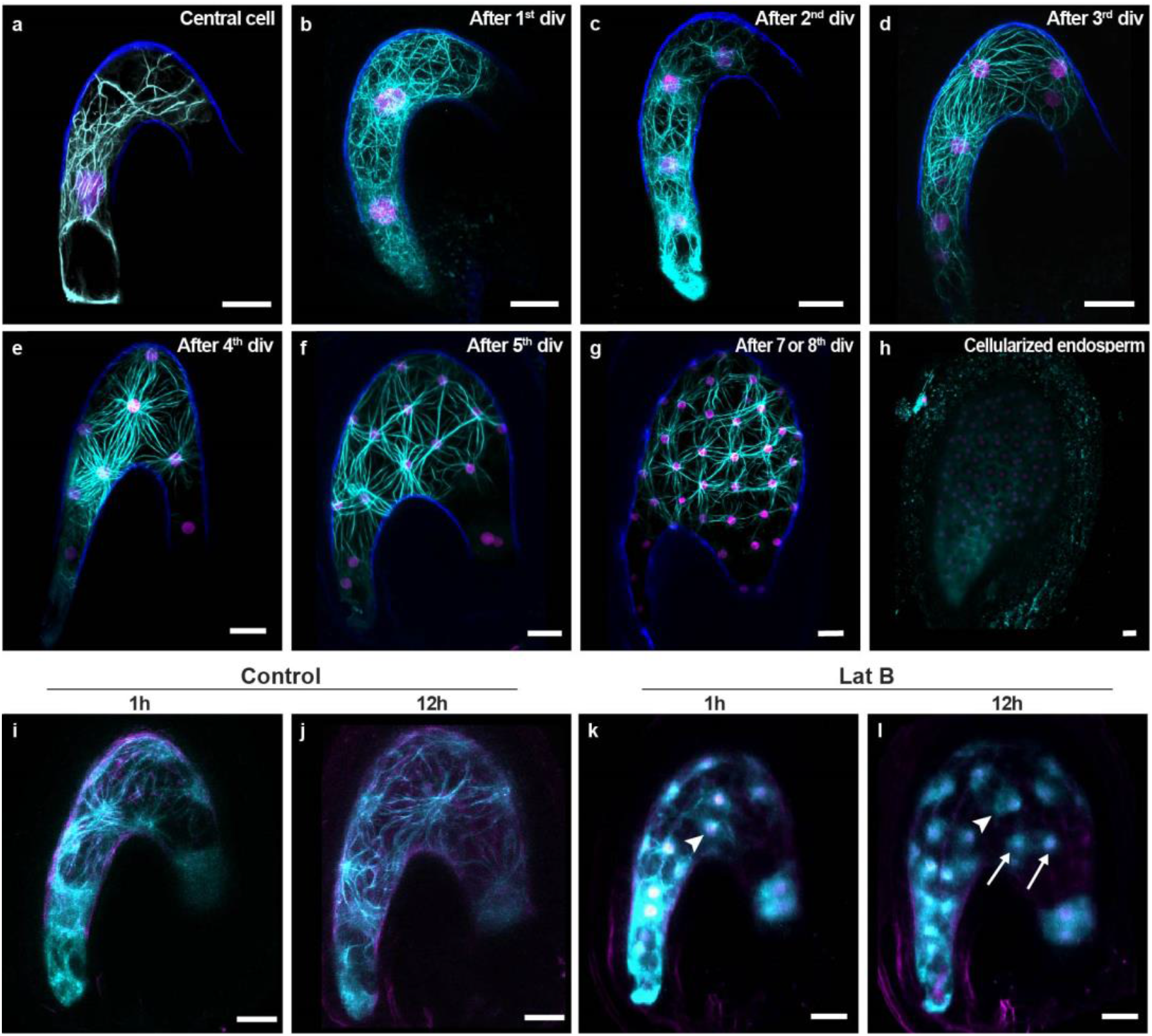
Unique aster-shaped structures of F-actin during coenocytic endosperm development. **a-l**, Z-projected confocal images of F-actin (cyan, *proFWA::Lifeact:Venus*), coenocytic endosperm nuclei (magenta, *proFWA::H2B:mRuby2*), and autofluorescence (blue) from the central cell (**a**), after the 1^st^ to 5^th^ divisions (**b-f**) and 7^th^ or 8^th^ division (**g**), and cellularized endosperm (**h**). Time-lapse Z-projected confocal images showing that Lat B treatment disrupted F-actin but did not inhibit nuclear divisions, control 1h and 12h after mock treatment (**i, j)** and Lat B 1h and 12h after treatment (**k, l**). Arrowheads indicate disrupted F-actin and the arrows indicate dividing nuclei. **f** corresponds to video 2, **i-l** correspond to video 4. Scale bar, 20 µm.

After the 3^rd^ nuclear divisions, the central vacuole develops and pushes the cytoplasm including nuclei to the plasma membrane periphery (Video 2). F-actin generated unique aster-shaped structures between each nucleus and the plasma membrane, and these were connected to each other through long filaments (Fig. 2d-g, Extended Data Fig. 2 and Video 2-3). F-actin asters were visible during the remaining coenocytic endosperm development (Fig. 2d-g and Video 3) and disappeared upon cellularization (Fig. 2h).

Treatment with the F-actin depolymerizing drug, Latrunculin B (Lat B), caused random bouncing-like movements of nuclei, especially immediately after nuclear divisions (Video 4). In the control, daughter nuclei moved away from the position of their mother nucleus and upon reaching the maximal displacement, they maintained their positions until the next round of nuclear division (Extended Data Fig. 3 and Video 4). By contrast, in the Lat B treatment, nuclei kept moving further after nuclear division, nearly colliding with neighboring nuclei and bouncing back and forth (Fig. 2i-l and Video 4). Consistently, the endosperm-specific expression of the semi-dominant negative *ACTIN* gene generating fragmented F-actin (*proFWA::DN-ACT8*, hereafter referred to as *DN*-*ACTIN*^19^) also showed the random bouncing-like movement of nuclei after nuclear division (Fig. 3a-c, e-g and Video 5). These data indicate that F-actin does not play a major role in pulling the daughter nuclei immediately after the division, but it restricts further movement and controls their equidistant positioning in the coenocytic PEN.

**Figure 3:**
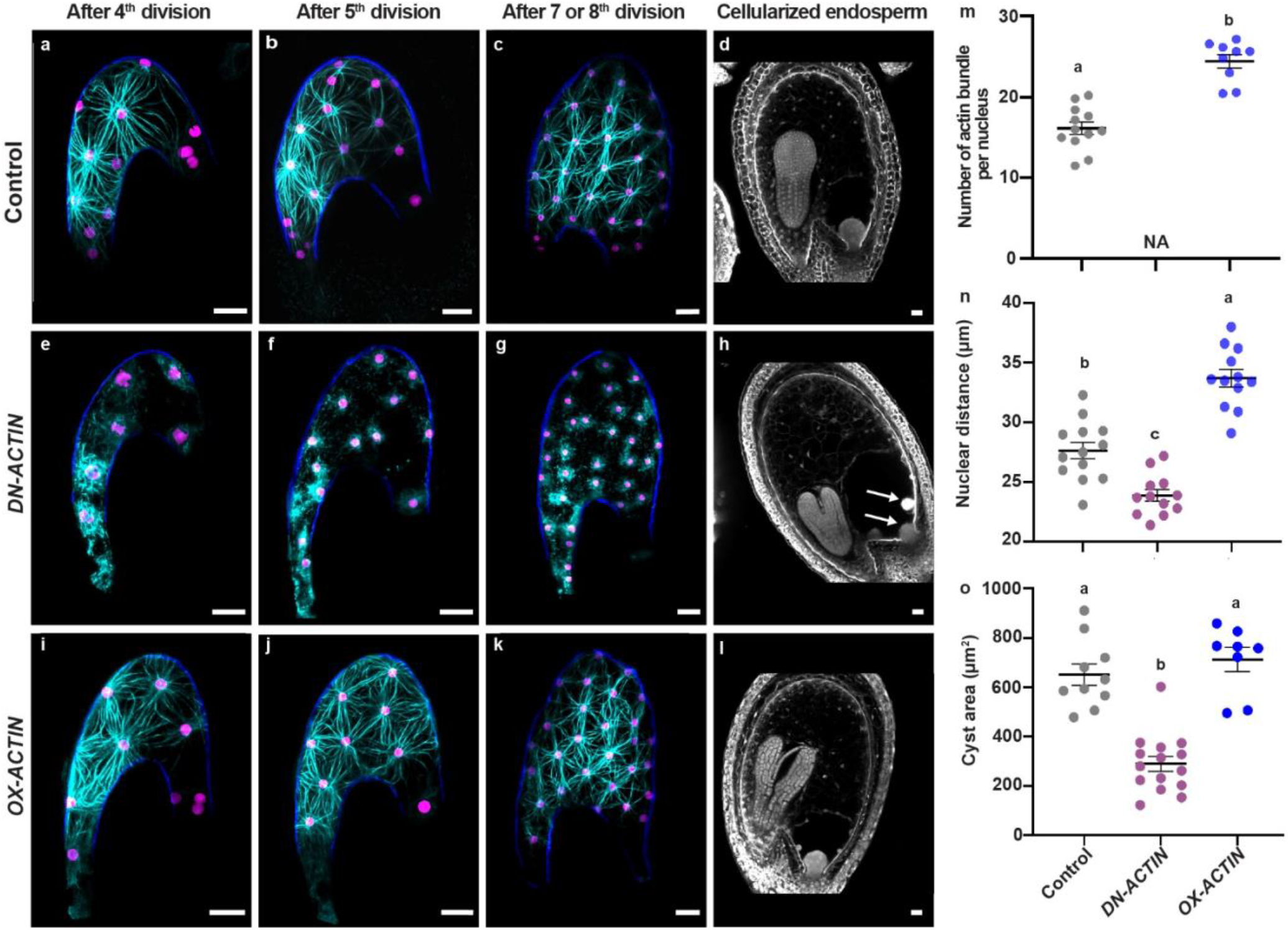
F-actin controls nuclear organization during coenocytic endosperm development. **a-c, e-g, i-k**, Z-projected confocal images of F-actin (cyan, *proFWA::Lifeact:Venus*), coenocytic endosperm nuclei (magenta, *proFWA::H2B:mRuby2*), and autofluorescence (blue) in the coenocytic endosperm. Control (**a-c**), dominant negative actin *(DN-ACTIN)* (**e-g**), and overexpressed actin *(OX-ACTIN)* (**i-k**). Scale bar, 20 µm. **d, h, l**, Z-projected confocal images of Feulgen-stained cellularized endosperm at 6 days after pollination (DAP), control (**d**), *DN-ACTIN* (**h**), *OX-ACTIN* (**l**). Arrows in **h** indicate multiple cysts. Scale bar, 20 µm. **m-n**, Quantitative analysis of actin bundles around each nucleus (**m)** and internuclear distance (**n**) in coenocytic endosperm after the 5^th^ division. Individual dots represent the means of actin bundle number and internuclear distance per seed, and black bars on the dot plots represent the mean of the means. Error bars represent the standard errors. NA denotes not applicable, because no actin bundle. **o**, Cyst size in the control, *DN-ACTIN*, and *OX-ACTIN*. Individual dots in the control and *OX-ACTIN* represent the area of a single cyst from a single seed, and dots in *DN-ACTIN* represent the areas of multiple cysts from a single seed (six seeds total). Levels not connected by the same letter (a–c on the graph **m-o**) are significantly different (*p* < 0.01, Tukey-Kramer HSD test).

In the CZE, Lifeact-Venus did not visualize any obvious structures in the CZE region where chalazal nodules are generated and moved to the chalazal cyst (Fig. 2e-g and Video 3). Despite the lack of characterization of F-actin in the CZE, we observed abnormal cyst formation in *DN*-*ACTIN* (Fig. 3h). In the control, one large cyst is present at the chalazal pole (Fig. 3d), whereas in *DN*-*ACTIN*, there were multiple small cysts that could not move towards the chalazal pole (Fig. 3d, h, o and Extended Data Fig. 1), indicative of a role for F-actin in depositing and incorporating nuclei at the chalazal pole. Lifeact recognizes F-actin by binding to a hydrophobic pocket on two adjacent actin subunits preferentially with the closed D-loop (DNase I binding loop), a hallmark of ADP states of F-actin^31,32^. The hydrophobic binding site of F-actin overlaps with the binding region of actin binding proteins, such as cofilin and myosin, resulting in binding competition between Lifeact and actin binding proteins^31^. The class XI myosin, *XIG* (*AT2G20290*) and cofilin (*AT3G45990*) are highly enriched in CZE^33,34^. It still remains unclear why F-actin is not visible in the CZE while playing a role in nuclear transport. Further analyses are awaited to reveal whether distinctive conformational changes of F-actin that might have altered Lifeact binding affinity^31^ and/or intensive competition of Lifeact with competitors such as *XIG* and cofilin exist in the CZE.

To further understand the role of F-actin in coenocytic endosperm development, we monitored the dynamics in an endosperm-specific *ACT8* over-expressing line (*proFWA::ACT8*, hereafter referred to as *OX*-*ACTIN*^35^; Fig. 3i-k). Over-expression of actin isoforms can change actin organization such as increasing actin bundling or density by massive polymerization likely due to the increased concentration of G-actin^36,37^. Compared to the control, *OX*-*ACTIN* showed a significantly increased number of actin bundles on each nucleus and longer internuclear distances (Fig. 3i-k, m, n). The overall structure of F-actin in the endosperm as well as the nuclear division pattern, intervals, movements, and formation of the cyst in *OX-ACTIN* were similar to those in the control (Fig. 3a-d, i-o, Extended Data Figs. 1, 4 and Video 6). These results further support the notion that it is not a delicate balance of actin dynamics, but rather the unique F-actin structures that are important in the arrangement and movement of endosperm nuclei during coenocytic endosperm development.

### MTs contribute to the foundation of F-actin organization and regulates nuclear movement

Coenocytic endosperm MT (*proFWA::TagRFP-TUA5*) asters radiated from each nucleus during mitosis (Fig. 4a-d, f and Video 7), consistent with the immunofluorescence patterns previously reported^21–24^. The dynamics of coenocytic endosperm MTs during mitosis and interphase was similar with that in somatic cells (Fig. 4e and Video 8)^38,39^. The nuclear-based MT asters became more apparent after the 3^rd^ nuclear divisions, coinciding with F-actin aster formation (Figs. 2 and 4b-d, Extended Data Fig. 2). F-actin asters radiating over the nucleus co-localized with MTs (Fig. 4g-k). When MTs formed spindles during the mitotic phase, F-actin asters became disorganized concomitantly and reassembled with the formation of MT asters after nuclear division (Extended Data Fig. 5). To further investigate the relationship between F-actin and MTs, we monitored the dynamics of MTs and F-actin in the presence of Lat B and oryzalin (an inhibitor of MT polymerization)^40^. With 10 μM oryzalin, the endosperm MT asters became less apparent within 3 hours of the treatment, nuclear division failed, and the nuclei aggregated (Fig. 5a-d, Extended Data Fig 6, and Video 8). Treatment with 20 μM oryzalin for 1 hour disrupted overall MTs, also resulting in failed nuclear division and seeds that then collapsed (Extended Data Fig. 6). F-actin organization was also affected by oryzalin; after most MT asters disappeared, the F-actin asters became a disorganized reticulate pattern uncentered on the nuclei (Fig. 5e-h and Extended Data Fig. 6 and Video 9). These results suggest that MT asters are required for F-actin aster formation in the coenocytic endosperm. To determine whether, and if so, when F-actin asters recover after MT aster reconstruction, we performed time-lapse F-actin imaging after the oryzalin wash-out (20 μM for 1 hour). Both MT and F-actin asters reappeared approximately 2 and 7 hours after washout, respectively, and normal mitotic nuclear divisions and movement followed (Fig. 5i-p, and Video 10). Lat B treatment did not affect the overall MTs organization (Extended Data Fig. 7, and Video 8), consistent with the result that nuclear division controlled primarily by MT was normal in both Lat B and *DN-ACTIN* (Figs. 2k-l and 3e-g and Videos 5 and 6). Taken together, these results indicate that the MT asters generated immediately after nuclear division serve as the foundation for the aster-shaped F-actin that restricts nuclear movement and controls nuclear equidistance.

**Figure 4:**
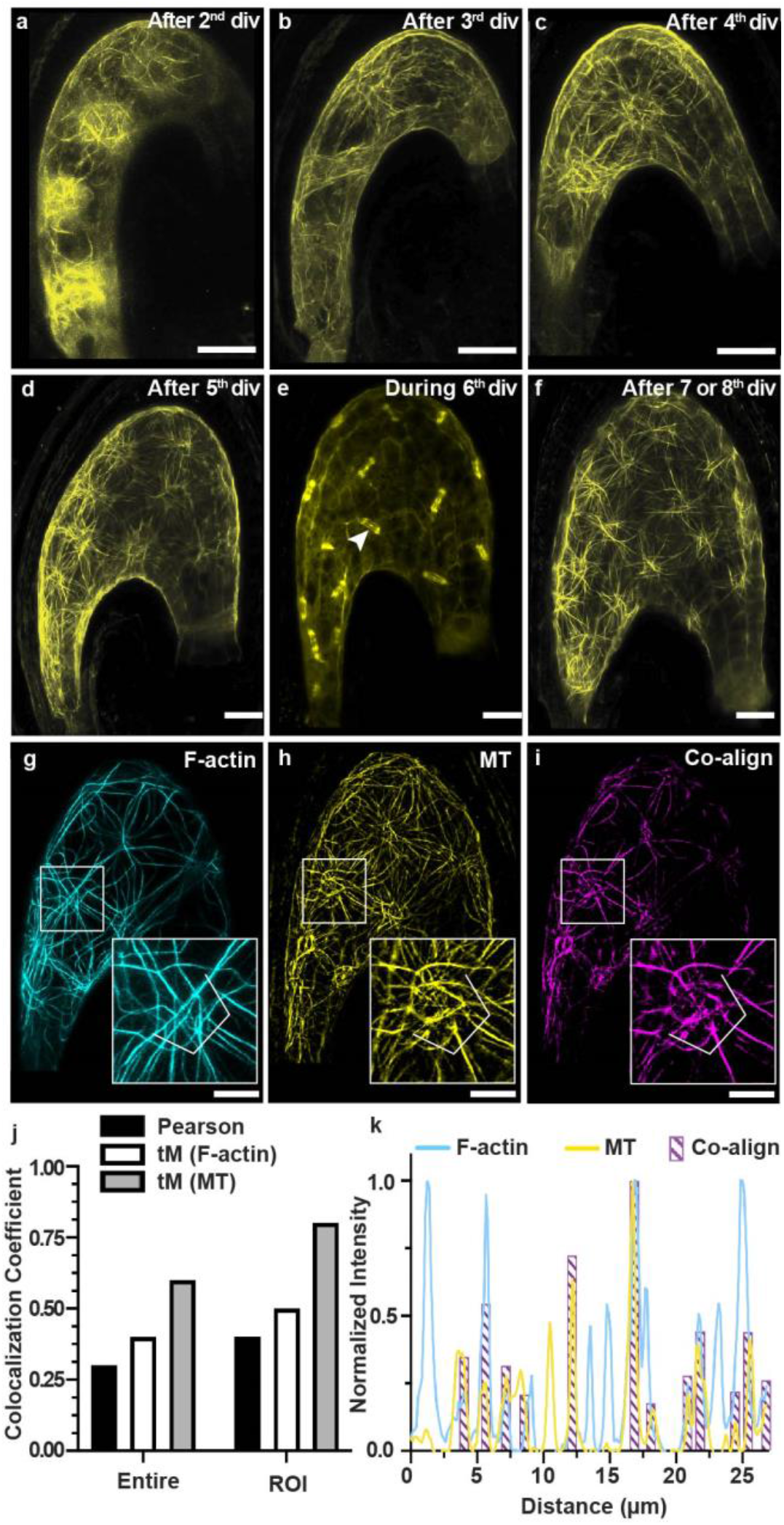
MT organization and crosstalk between F-actin and MTs in coenocytic endosperm. **a-f**, Z-projected confocal images of coenocytic endosperm MTs (yellow, *proFWA::TagRFP::TUA5*) after the 2^nd^ to 5^th^ (**a**-**d**) and 7^th^ or 8^th^ divisions (**f**). MT spindle formation during mitotic phases (**e**). The white arrowhead in **e** indicates MT spindle formation during mitotic nuclear division. **g-n**, Co-localization of F-actin and MTs in the coenocytic endosperm. Background-subtracted two channel images of F-actin (**g**, cyan, *proFWA::Lifeact:Venus*) and MTs (**h**, yellow, *proFWA::TagRFP::TUA5*) from a double marker line of F-actin and MT, and co-alignment of F-actin and MT (**i**, magenta). Scale bar, 20 µm. The region of interest (ROI, white square line in **g-i**) including a single nucleus used for the following co-localization analysis (**j-k**) was enlarged and inserted in the right bottom of each image (**g-i**). **j**, Quantification of co-localization by calculating the Pearson’s coefficient (black bar) and the thresholded Manders’ (tM) coefficient per each channel (white bar, F-actin; gray bar, MTs). The co-localization coefficient values were calculated from entire endosperm (left) and the ROI (right) on the graph. **k**, Normalized intensity profiles of F-actin (cyan), MT (yellow) and the co-alignment (magenta) around the nucleus (white line) in the ROI **(g-i)**. Note that F-actin, which does not show co-alignment with MTs at this position, co-aligns with MTs at different positions (depth).

**Figure 5:**
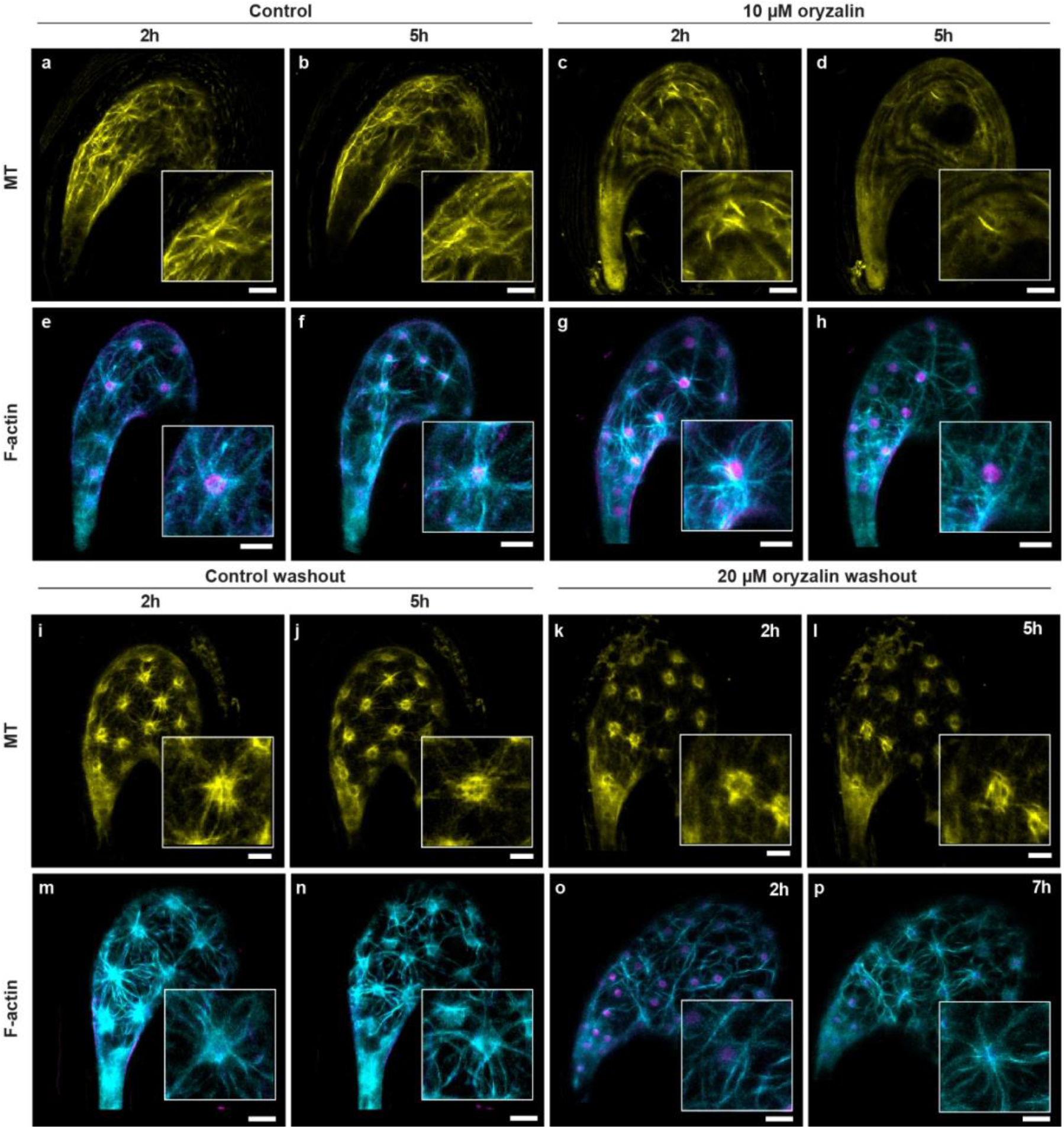
MT asters are required for the organization of F-actin aster structures. **a-h**, Time-lapse Z-projected confocal images of coenocytic endosperm MT (yellow, *proFWA::TagRFP::TUA5*) and F-actin (cyan, *proFWA::Lifeact:Venus*) with endosperm nuclei (magenta, *proFWA::H2B:mRuby2*) after mock and oryzalin treatments. MT control 2 h and 5 h after mock treatment (**a, b**), MT 2 h and 5 h after 10 µM oryzalin treatment (**c, d**), F-actin control 2 h and 5 h after mock treatment (**e, f**), and F-actin 2 h and 5 h after 10 µM oryzalin treatment (**g, h**). **a-d** correspond to Video 8, and **e-h** correspond to Video 9. **i-p**, Time-lapse Z-projected confocal images of coenocytic endosperm MTs and F-actin after washout of 20 µM oryzalin. MTs 2 h and 5 h control washout (**i-j**), MT 2 h and 5 h after oryzalin washout (**k, l**), F-actin 2 h and 5 h control washout (**m, n**), and F-actin 2 h and 7 h after oryzalin washout (**o, p**). Enlarged inserts at the bottom right of each image represent the F-actin and MTs around the nucleus. **i-p** correspond to Video 10. Scale bar, 20 µm.

**Figure 6:**
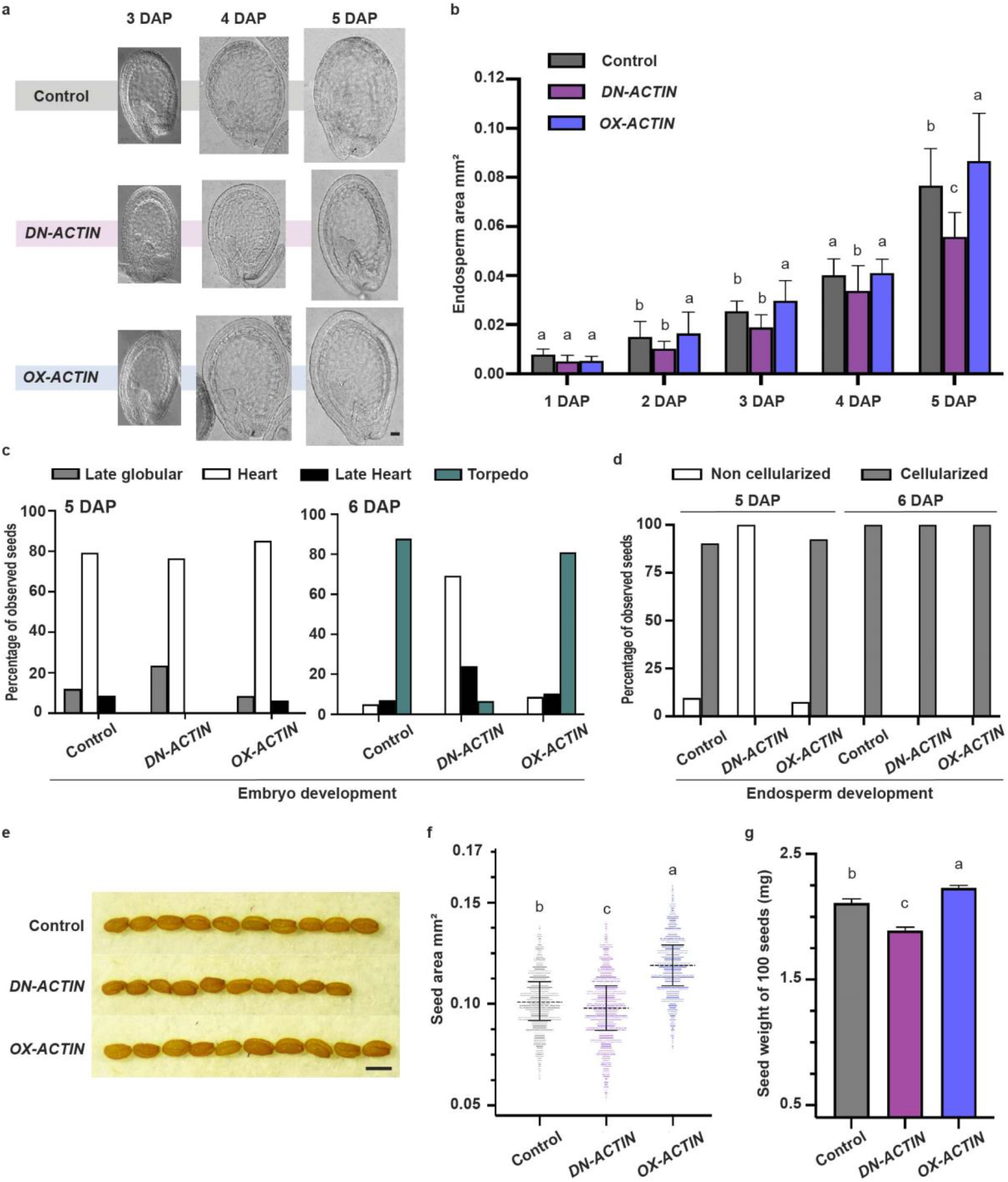
F-actin dynamics in the coenocytic endosperm affect the endosperm and final seed size. **a**, DIC microscopy of cleared whole-mount control, *DN-ACTIN*, and *OX-ACTIN* seeds at 3, 4, and 5 DAP. Scale bar, 50 µm. **b**, Average area of the coenocytic endosperm in control, *DN-ACTIN*, and *OX-ACTIN* seeds from 1 to 5 DAP. 1-2 DAP, n=10 seeds; 3-5 DAP, n=15-20 seeds. Error bars represent the standard error. Endosperm areas at the same stage were compared statistically. **c-d**, The embryo (**c**) and endosperm (**d**) developmental stages observed in the control, *DN-ACTIN*, and *OX-ACTIN* seeds at 5 and 6 DAP by Feulgen staining analysis. 5 DAP Control, n= 104; *DN-ACTIN*, n= 64; *OX-ACTIN*, n= 176; 6 DAP Control, n= 140; *DN-ACTIN*, n= 165; *OX-*h*ACTIN*, n= 173. **e**, Comparison of mature seeds of the control, *DN-ACTIN* and *OX-ACTIN*. Scale bar, 500 µm. **f-g**, Quantitative analysis of seed size (**f**) and 100-seed weight (**g**) of mature seeds in the control, *DN-ACTIN*, and *OX-ACTIN*. The seed size of each line is represented by 1,000 seeds per plant from six individual plants. The middle black dotted line within the plot shows the median. Seed weight is the average value of 10 sample batches each containing 100 seeds. Error bars represent the standard error. Levels not connected by the same letter (a–c on the graph **b, f, g**) are significantly different (*p* < 0.01, Tukey-Kramer HSD test).

### F-actin dynamics in the coenocytic endosperm affect the endosperm and final seed size

The size of *DN-ACTIN* coenocytic endosperm remained smaller with shorter internuclear distances compared to the control (Figs. 3n and 6a, b). While the endosperm nuclei divisions at the beginning were not significantly altered in *DN-ACTIN*, the division intervals in later stages from the 6^th^ division became longer than those of the control and the cellularization was delayed (Extended Data Fig. 4). In the control, 80% of the seeds reached the heart-shaped embryo stage and the endosperm started cellularizing at 5 DAP, and at 6 DAP, 85% of the embryos reached the torpedo-shaped embryo stage (Fig. 6c, d and Extended Data Fig. 1). Whereas in *DN-ACTIN*, 77% of the embryos reached the heart-shaped stage without endosperm cellularization at 5 DAP, and just 7% of the embryos reached the torpedo-shaped stage at 6 DAP (Fig. 6c, d and Extended Data Fig. 1). The timing of cellularization is known to be highly associated with the seed size^8,13,14^; however, *DN-ACTIN* produced smaller seeds with a longer coenocytic endosperm phase compared to the control (Fig. 6e-g).

In contrast to *DN-ACTIN*, we observed larger seeds in *OX-ACTIN* compared to the control (Fig. 6e-g). *OX-ACTIN* did not show any change in nuclear division numbers and intervals in the coenocytic endosperm (Extended Data Fig. 4) but caused enlarged coenocytic endosperm (Fig. 6a, b). The *OX-ACTIN* coenocytic endosperm showed an increased number of actin bundles on each nucleus with longer internuclear distance compared to the control (Fig. 3i-k, m, n). The developmental speed of both embryo and coenocytic endosperm did not differ in the control and *OX-ACTIN* (Fig. 6c, d and Extended Data Fig. 4). Taken together, these results show that the coenocytic endosperm area before cellularization in these endosperm-actin-manipulated lines reflects the mature seed size and there is no positive correlation between the duration of the coenocytic endosperm development and the final seed size (Fig. 6b, f and Extended Data Fig. 4).

## Discussion

This work has revealed the details of coenocytic endosperm dynamics that highlight the unique function of F-actin for the organization of endosperm nuclei, the requirement of MTs for F-actin aster organization, and the role of F-actin in seed size determination. The *Drosophila melanogaster* embryo has been intensively studied as a coenocyte model where both F-actin and MTs actively control coenocyte nuclear dynamics^41,42^. Similar to *Arabidopsis*, MTs generate the force to pull daughter nuclei apart during mitotic phase and F-actin restricts the movement and maintains the position of these daughter nuclei in the *Drosophila* coenocytic embryo^41,42^. However, while MTs show similar nucleus-centered aster structures at interphase in both species, *Drosophila* F-actin displays a dome-like accumulation between the plasma membrane and each nucleus at interphase in the actin-rich cortex^41–45^. *Arabidopsis*, on the other hand, generates F-actin asters between the plasma membrane and each nucleus (Video 2), and the difference in F-actin structure between them is possibly due to an additional function of F-actin in *Drosophila*. The dome-shaped F-actin acts as an anchoring platform to hold nuclei at the cortex^41,43,45^. In *Arabidopsis*, there is a large central vacuole in the coenocytic endosperm, pushing the cytoplasm to the plasma membrane periphery and the endosperm nuclei do not need any active anchoring system to maintain their positions close to the plasma membrane periphery (Video 2). F-actin in *Arabidopsis* might simply be reassembled by MT, stay co-aligned with MT as asters, and play a role in restricting nuclear movement after division and controlling the distances among nuclei in PEN and MCE. In the CZE, PEN nuclei were pulled or pushed by F-actin towards the chalazal pole to generate the cyst (Video 3). In *DN-ACTIN*, the deposition of nuclei in the chalazal pole was disturbed, resulting in multiple small-sized cysts in the CZE (Fig. 3h, o). These results suggest that in addition to restricting the nuclear movement in PEN, F-actin has a role in nuclei movement toward the chalazal pole. The formin *AtFH5*, one of the actin nucleators, is highly expressed in the chalazal endosperm and the mutant shows smaller cysts or absence of cyst formation^46^, further supporting the involvement of F-actin in CZE nuclei deposition during cyst formation.

After double fertilization in *Arabidopsis*, rapid proliferation of the coenocytic endosperm through mitosis without cytokinesis governs the increase in seed volume until endosperm cellularization occurs. Precocious endosperm cellularization can result in relatively smaller seeds, while delayed endosperm cellularization is associated with enlarged seeds in *Arabidopsis* and rice^8,13,14^. In addition, because of the potential importance of the cyst for maternal nutrient transfer to the seed^47,48^ as well as enlargement of the cyst in the larger seeds of the Polycomb Repressive Complex 2 mutants (PRC2)^49^, the cyst has been considered to be linked with the final seed size. However, in our experiments, delayed endosperm cellularization with the small cyst was observed in *DN-ACTIN* that produced smaller seeds, and *OX-ACTIN* showed no change in either endosperm cellularization timing or cyst size, yet it produced larger seeds (Figs. 3o, 6d-f). Plants carrying double mutations in the PRC2 pathway and HAIKU pathway generate smaller seeds with the enlarged cyst^16^, also supporting the idea that the cyst size is not linked with the final seed size. The cyst enlargement in the PRC2 mutants is likely caused by the continuous incorporation of PEN nuclei due to the absence of endosperm cellularization.

What then, can cause the size increase in the coenocytic endosperm and the final seed in *OX-ACTIN*? We did not observe any changes in the dynamics of F-actin or nuclei between the control and *OX-ACTIN* (Fig. 3). However, *OX-ACTIN* showed more actin cables and bundles with longer internuclear distance, whereas *DN-ACTIN* exhibited shorter internuclear distances and smaller sizes of the coenocytic endosperm and final seed (Figs. 3m, n and 6b, e-g). Overexpression of *ACTIN* genes in somatic cells does not generate enlarged plants^37^, but in the coenocytic endosperm, the additional F-actin could allow tethering of nuclei further apart. Possibly the longer actin cables provide the force to expand the endosperm cell. Another possibility is that F-actin controls turgor pressure in the coenocytic endosperm. Together with MTs, F-actin plays a key role in determining plant cell shape, mainly by affecting all modes of cell expansion, which is tightly linked with turgor pressure^50–52^. Endosperm derived-turgor pressure of the seed is maximized at the coenocytic stage and later constrained by cellularization^52^. It is possible that F-actin contributes to the control of the endosperm turgor pressure or targeting regulators of cell wall properties, eventually contributing to the regulation of endosperm expansion before cellularization, which pre-determines the final seed size^16^. Insights into the role of F-actin in the size of the coenocytic endosperm as well as determination of the final seed size are elucidated in this study, providing new targets for strategies to increase seed size for our food security.

## Methods

### Plant material and growth condition

All plant lines: the F-actin control (*proFWA::Lifeact:Venus;proFWA::H2B:mRuby2*), *OX-ACTIN* (*proFWA::Lifeact:Venus;proFWA::H2B:mRuby2;proFWA::ACT8*), *DN-ACTIN* (*proFWA::Lifeact:Venus;proFWA::H2B:mRuby2;proFWA::DN-ACT8)*, and MT (*proFWA::TagRFP-TUA5*) marker lines used in this work were all derived from the *Arabidopsis thaliana* Columbia-0 (Col-0) ecotype. Seeds were germinated and the seedlings were grown for two weeks under short-day conditions (8 h light, 22°C and 16 h dark, 18°C). Plants were then grown with continuous light at 22°C. The constructs *proFWA::Lifeact:Venus, proFWA::H2B:mRuby2, proFWA::ACT8*, and *proFWA::DN-ACT8* have been described previously^18,19,35^. The transgenic line carrying *proFWA::Lifeact:Venus* was crossed with the MT marker line, *proFWA::TagRFP-TUA5* to generate a double marker line of F-actin and MTs.

### Plasmid construction and transformation

The DNA construct used in the MT marker line was generated using Multisite Gateway Technology (Invitrogen, CA, USA). The multisite gateway binary vector pAlligatorG43 and entry clones of pENTRP4P1r-proFWA and pENTR221-TagRFP-TUA5, described previously^20^, were recombined into pAlligatorG43 to generate *proFWA::TagRFP-TUA5* and transformed into *Arabidopsis* Col-0 using the floral dip method^53^.

### Sampling for the live-cell imaging and chemical preparation

*Arabidopsis* siliques were dissected with a sharp knife and developing seeds were collected into an assay medium (2.1 g/L Nitsch basal salt mixture, 5% w/v trehalose dehydrate, 0.05% w/v MES KOH (pH 5.8), and 1X Gamborg vitamins) in a multi-well glass-bottom dish as described previously^54^. For each experiment, seeds from 4-5 siliques were collected into 200-µL assay medium. For long live-cell imaging, 0.5% low-melting agarose and 0.1-µL Plant tissue culture contamination control (P6820, Phyto Technology Laboratories) were added to the 200-µL assay medium. To observe division of the primary endosperm nucleus, pistils were pollinated 12h before sample collection. Lat B (stock, 10 mM; Sigma-Aldrich, MO, USA) and oryzalin (stock, 10 mM; Sigma-Aldrich, MO, USA) stock solutions were prepared in DMSO (dimethyl sulfoxide) and kept at −80°C. Freshly prepared working concentrations of Lat B (5 µM) and oryzalin (10 µM and 20 µM) were prepared before each experiment in the assay buffer. To remove oryzalin, seeds were washed 3-4 times at 10 min intervals with the assay medium.

### Confocal microscopy and image processing

All time-lapse confocal images were captured using a FV1200 laser scanning confocal microscope system (Olympus) equipped with 515-nm, and 559-nm lasers. A GaAsP detection filter was used to detect Lifeact:Venus (Ex 515-nm), H2B:mRuby2 (Ex 559-nm) and tagRFP:TUA5 (Ex 559-nm). All time-lapse images were acquired with a 40X dry objective lens. Time-lapse (15-30 min interval) images with z-planes (25-35 µm total, 3-4 µm each slice) were acquired using FV10-ASW 4.2 software. Laser 3-4%, HV 500-550, gain 1.25 and Kalman 2 options were used to capture images. All Z-projected static confocal images were captured using an FV3000 laser scanning confocal system (Olympus) equipped with 514-nm and 561-nm laser lines. Z-projected confocal images were acquired with a 30X silicon oil immersion objective lens. The confocal images with z-planes (30-40 µm total, 0.76 µm each slice) were acquired using FV31S-SW software. Laser 2-3%, HV 500-550, and gain 1.25 options were used to capture images. All images were processed by constrained iterative deconvolution using CellSens Dimension Desktop 3.2 (Olympus) to improve quality. Autofluorescence detected from the RFP channel (Em 560-620 nm) was removed by Imaris 9.7.2 (BitPlane) spot detection and masking options. Images obtained from the F-actin and MT double marker line were background-subtracted using Fiji (Image J) in both YFP (Em 500-560 nm) for F-actin and RFP (Em 560-620 nm) for the MT channels to remove background noise. The co-localization Pearson’s and thresholded Manders’ coefficient values of the YFP (F-actin) and RFP (MT) channels from the entire endosperm and region of interest (ROI; Fig. 4g-i) were analyzed using Imaris Coloc function and then a co-alignment channel was created. The localization of F-actin and MT and their co-alignment were shown in different pseudo-colors using Fiji. For histogram profiling of the YFP (F-actin), RFP (MT), and the co-alignment channels from the ROI, the lines of analysis points were drawn and the intensity plots along the lines from each channel were obtained using Fiji. The obtained intensities were normalized by dividing by the maximum value of each channel.

### Nuclei division interval measurement

To measure nuclei division intervals, seeds were first categorized based on the nuclei number and size of the seeds we observed at the start time of time-lapse imaging. The time from the start of imaging to the initial nuclei division was not used for interval measurement. The measured division interval times from the subsequent divisions of multiple seeds were aligned based on the division stage and averaged.

### Nuclear movement measurement

The coenocytic endosperm after the 5^th^ division in the control, *OX-ACTIN* and *DN-ACTIN* were imaged by confocal microscopy and 3D images were created by Imaris 9.7.2 (BitPlane). All endosperm nuclei were outlined as spheres using Imaris spot detection and were used to calculate the internuclear distances. To determine the internuclear distances in the control and *OX-ACTIN* lines, the center nucleus of the PEN and the neighboring nuclei along with the F-actin asters were selected. The distance between each of the neighboring nuclei and the center nucleus was then measured. Because both the F-actin and nuclear position were disrupted in *DN-ACTIN* (Fig. 3e-g), the distances between the nuclei in the periphery were measured, not the nuclei located close to other nuclei. The multiple distances from one seed sample were averaged. To measure nuclei displacement, Z-projected confocal images were processed in Fiji (ImageJ) using the tracking function from the Manual tracking plugin. From the onset of nuclear division until 2h, the daughter nuclei were tracked manually to record nuclear displacement.

### Number of actin bundles measurement

The coenocytic endosperm after the 5^th^ division in the control, *OX-ACTIN*, and *DN-ACTIN* were imaged by confocal microscopy and 3D structures were created by Imaris 9.7.2 (BitPlane). In each 3D image, 5-7 PEN nuclei were selected to count the F-actin bundles manually. The total number of bundles from 5-7 selected nuclei was then averaged. For each line, the mean and standard error of the averages from 10 to 12 images were shown.

### Orientation analysis

To analyze the distribution of F-actin and MT aster structures surrounding the endosperm nucleus, a square ROI where the nucleus is center was set in each image and the frequency histogram of spatial orientations of F-actin and MT was calculated using the OrientationJ Distribution plugin (http://bigwww.epfl.ch/demo/orientation/) of Fiji. To evaluate aster structures, the cables above the nucleus were excluded from the calculation by masking the nucleus on F-actin and MT channel images, respectively. The ROI was quartered by four squares on the basis of the nucleus center and the four squares were rotated to get the axis of a quarter nucleus used as a standard 0 degree (Extended Data Fig. 2). All frequency histograms of the orientations of F-actin and MTs relative to the axis were added from the four squares in the ROI and the frequency histogram was given as probability.

### Differential interference contrast (DIC) microscopy for endosperm area measurement

Siliques harvested at 1-5 DAP were opened on one side immediately after harvesting, transferred to fixing solution (ethanol (EtOH): acetic acid, 9:1) and stored overnight at 4°C. Siliques were carefully washed with 90% EtOH for 10 min followed by 70% EtOH for 10 min. They were then stored at 4°C in 70% EtOH until the next steps. The EtOH was removed, clearing solution (66.7% chloralhydrate, 25% H_2_O, and 8.3% glycerol) was added and incubated for 24 hours. After incubation, the valves and septum were removed and only the seeds were mounted on the slide with clearing solution. Cleared seeds were visualized by DIC using a Leica DM2500 LED microscope under 20X dry or 40X oil lenses. Endosperm area was determined manually by hand drawing in Fiji (ImageJ). Area values were obtained from Fiji (ImageJ) using the “Analyze particles” function. The means of 10-20 seeds from each sample were used for statistical analysis.

### Feulgen staining for determining endosperm cellularization

Seeds were prepared using the Feulgen staining method described previously^55^. In brief, siliques harvested 5-6 DAP were opened on one side with needles and were incubated in a fixing solution (EtOH:acetic acid 3:1). After 24 hours of incubation, the fixing solution was then replaced with 70% EtOH and the siliques were stored in 70% EtOH at 4°C until the next step. Stored samples were washed with water three times at 15 min intervals, incubated 1 h in 5N HCl followed by three washes with water at 15 min intervals. After that, samples were incubated in Schiff’s reagent for 4 h and then washed three times in cold water at 15 min intervals. Samples were again washed with 70% EtOH and then 95% EtOH at 10 min intervals, and were then washed 4-5 times with 99.5% at 5 min intervals. Samples were incubated again for 1h in EtOH:LR white resin (1:1) followed by overnight incubation in LR white resin only. After incubation, seeds were mounted on glass slides and baked with LR white at 60°C for 8h. Seeds were observed under an FV3000 laser scanning confocal system (Olympus) equipped with a 561-nm laser line with an excitation wavelength at 561 nm (Em 560-610). To get the cyst area, the cyst was marked manually by ImageJ on 6 DAP Feulgen staining endosperm images. Area values were obtained from Fiji (ImageJ) by the Analyze particles function.

### Seed size and weight measurements

Six plants from each line were grown together under the same conditions, and mature seeds were collected to obtain seed size and weight data. Seed size was measured as described previously^56^. In brief, 1,500-3,000 seeds were spread on a Perspex box and scanned using an EPSON V800. Scanned images were used to determine seed area from Fiji (ImageJ) by the following sequence of actions: Image > Adjust > Color threshold > Analyze > Analyze particles. To avoid dust and aggregated seeds, only particles 0.02 – 0.2 mm^2^ were measured. For each line, the seed size areas of 6,000 seeds, 1,000 from each plant, were analyzed using JMP pro16 software. To get the exact seed weight, ten sample batches each containing 100 seeds were counted and weighed manually.

### Statistics

All Tukey-Kramer HSD tests were performed using JMP pro16 software. All plots were prepared using GraphPad Prism 7 software.

## Supporting information

Supplementary Table and Figures and Vide legends

## Acknowledgments

We thank Drs. Anthony Clark and David Zaitlin for their critical comments on this manuscript and Dr. Yukinosuke Ohnishi for endosperm area analysis. This work was supported by NSF Grant IOS-1928836 (to T.K.); National Institute of Food and Agriculture, US Department of Agriculture Hatch Program Grant 1014280 (to T.K.).

