## Supplementary Table and Figures and Vide legends for "Cellular dynamics of endosperm development in *Arabidopsis thaliana*"

**Extended Data Table 1: Developmental stages of coenocytic endosperm.**

| Endosperm development stage | No. of endosperm nuclei | Interval of endosperm division | Description |
|---|---|---|---|
| 1st | 2 | 2h | Primary endosperm nucleus divides into two nuclei.<br>Two nuclei migrate to opposite end of primary endosperm.<br>F-actin and microtubule (MT) remain coiled around each nucleus. |
| 2nd | 4 | 3h | Two nuclei divide into four nuclei. |
| 3rd | 8 | 4.3h | Four nuclei divide into eight nuclei.<br>One of eight nuclei migrates close to the chalazal pole, divides and two nuclei remain there.<br>F-actin and MT organize asters around each nucleus.<br>From this division onwards, F-actin asters disappear during mitotic nuclear division, and appear again after division. |
| 4th | 16 | 6.3h | Eight nuclei divide into sixteen nuclei.<br>All endosperm compartments are established (MCE, PEN, and CZE).<br>From this division onwards, F-actin asters appear around each nucleus in MCE and PEN but not CZE. |
| 5th | 28 | 9.7h | Synchronous division of endosperm nuclei stops after 4th division.<br>Four nuclei at the chalazal pole do not divide at the 5th division.<br>These nuclei start to form the chalazal cyst. |
| 6th | Not determined | 12.2h | 6th to 9th nucler divisions share similar features as below:<br>Rapid expansion of endosperm occurs.<br>Few nuclei from the peripheral region migrate to the chalazal region to add to the chalazal cyst. |
| 7th | Not determined | 15.3h | |
| 8th | Not determined | 13.7h | |
| 9th | Not determined | 17.3h | |
| 10th | Not determined | 11.7h | 60 to 100 min after last division, endosperm starts to cellularize from the micropylar end.<br>After endosperm cellularization, the F-actin disappears from the endosperm |

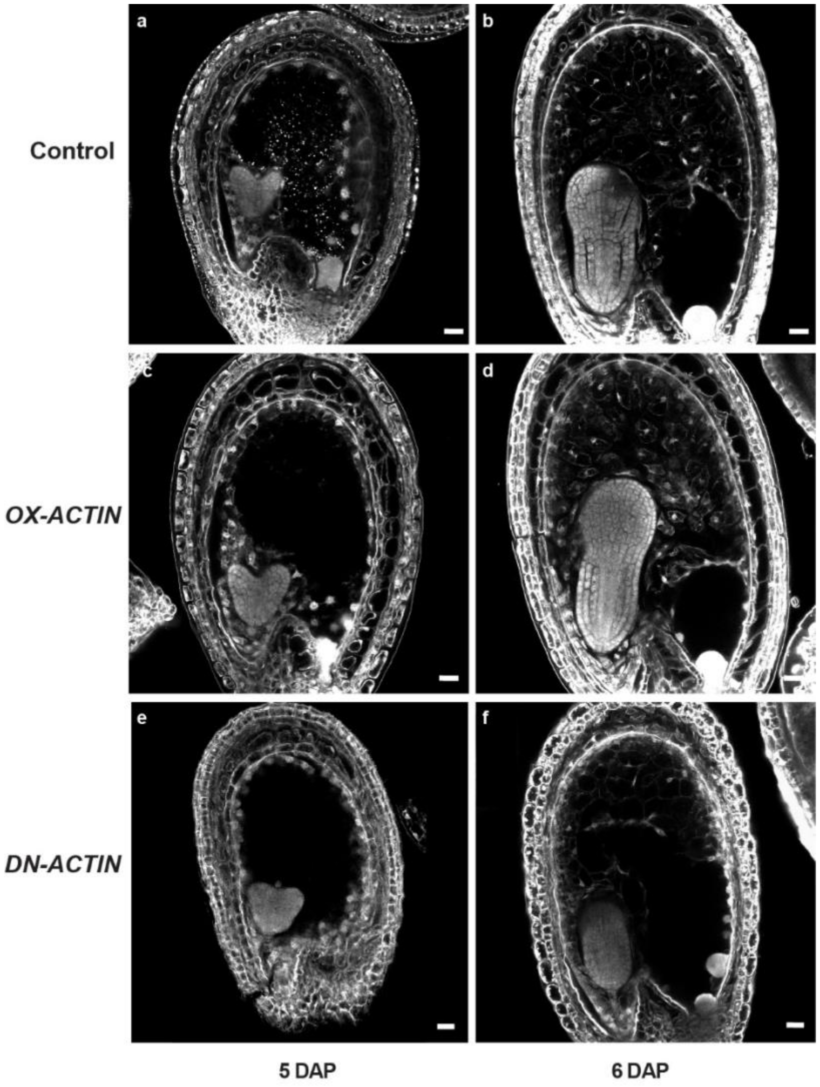


**Extended Data Figure 1: F-actin does not affect endosperm cellularization.** Z-projected confocal images of Feulgen-stained endosperm at 5-6 days after pollination (DAP). Control (**a-b**), *OX-ACTIN* (**c-d**), and *DN-ACTIN* (**e-f**). Endosperm cellularization was already observed in the control and *OX-ACTIN* at 5 DAP, whereas no endosperm cellularization has initiated in *DN-ACTIN*. Scale bar, 20 µm.

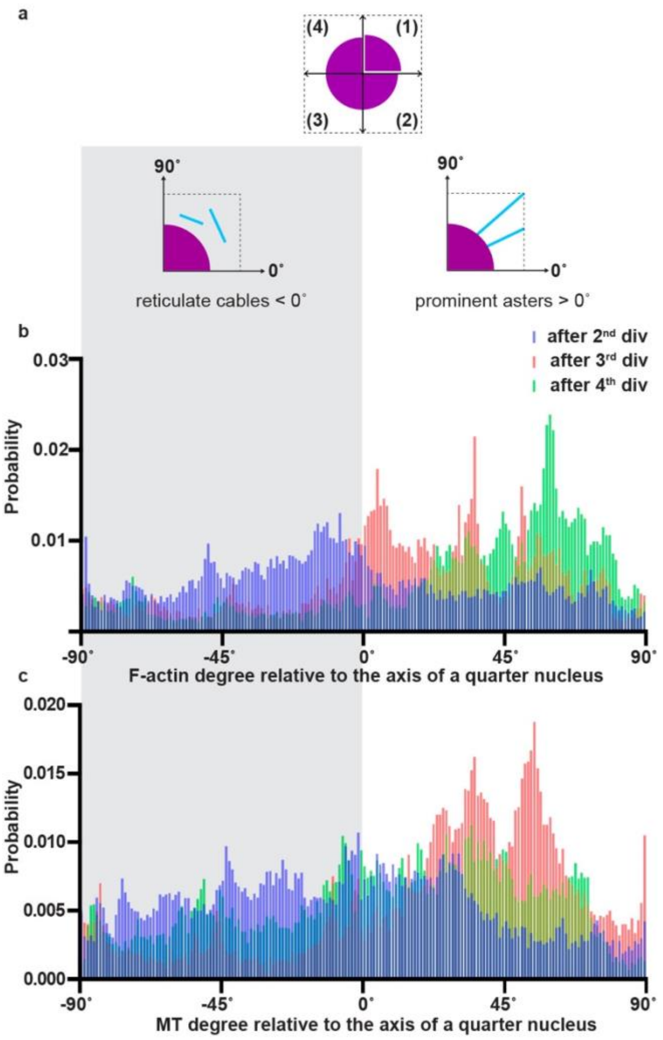


**Extended Data Figure 2: F-actin and MT aster structures become more apparent after the 3rd nuclear division. a,** Illustration indicating how to set the axis for analyzing frequency histogram of spatial orientations of F-actin and MTs surrounding endosperm nucleus (related to Figures 2c-e and 4a-c) and definition of the criteria to evaluate aster structure. Black square dot outline, ROI; magenta solid shapes, masked nucleus area; cyan solid lines, F-actin or MTs. The frequency histogram is given as probability distribution in the following graphs (**b-c**). **b-c,** Probability distribution of F-actin (**b**) and MTs (**c**) in degrees relative to the axis of a quarter nucleus (bottom in **a**). The probability is calculated from all frequency histograms of the orientations in F-actin and MTs combined from entire nucleus (top in **a**). Different colors denote different nuclear division stages. The shade indicates reticulate patterns of F-actin and MTs, which do not contribute to the aster structures.



43

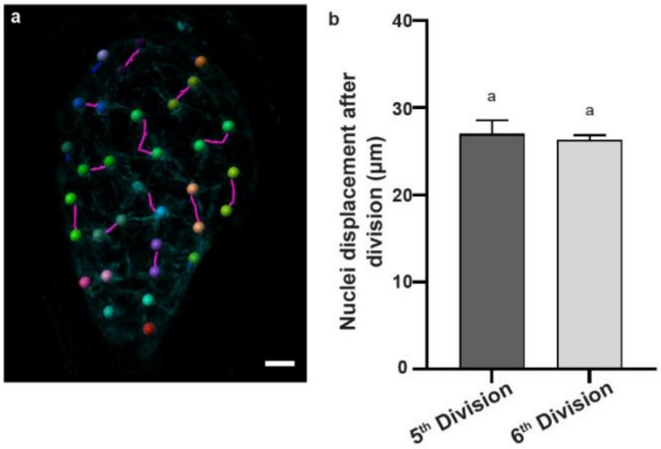


**Extended Data Figure 3: The displacement of daughter nuclei after mitotic nuclear division.** **a,** Z-projected confocal image of coenocytic endosperm nuclei (marked by different colored ovals) and their movement 2h after division. Nuclear trajectories of daughter nuclei after division are drawn by magenta. Scale bar 20 µm. **b**, Average nuclear displacement between 5th and 6th nuclear divisions. 5th division, n= 4 seeds (total 17 nuclei); 6th division, n=5 seeds (total 20 nuclei). Levels not connected by the same letter on the graph **b** are significantly different ($P < 0.01$, Tukey-Kramer HSD test). Error bars indicate the standard error.

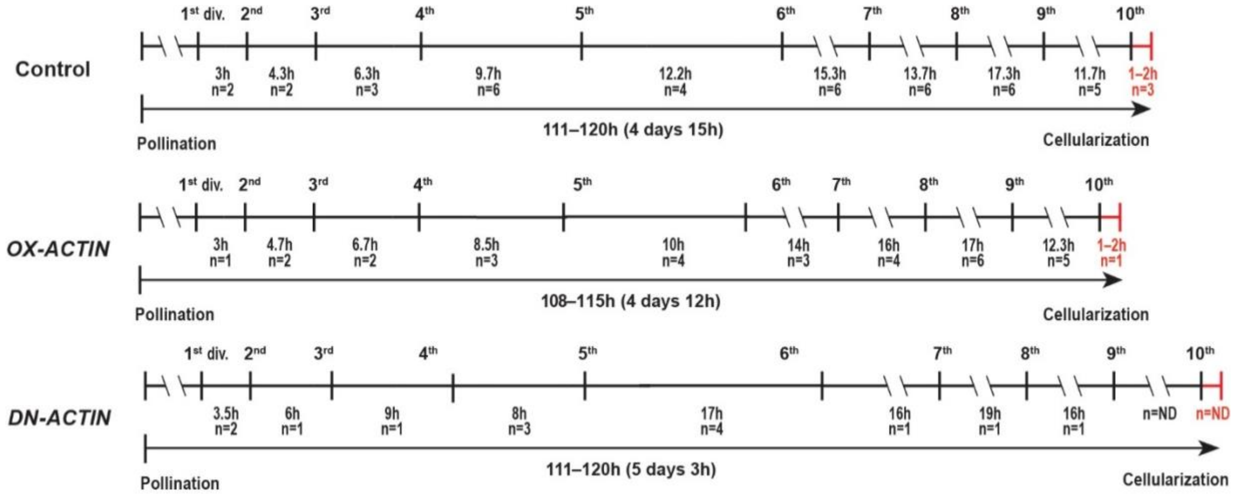


**Extended Data Figure 4: F-actin does not control nuclear division and intervals.** Coenocytic endosperm nuclear division numbers and intervals of control (top), *OX-ACTIN* (middle), and *DN-ACTIN* (bottom). Horizontal arrow bars represent the duration from pollination until endosperm cellularization and the duration represents the minimum and maximum of the total nuclear division intervals of individual samples in each line. Horizontal lines with vertical bars represent the average of each nuclear division interval. Distances between vertical bars are drawn to scale. In *DN-ACTIN*, we could not observe 9th and 10th division intervals due to the death of samples in long time-lapse live-cell imaging condition; however, the total duration was estimated based on the cellularization timing analyzed by Feulgen staining analysis (Fig. 6d). ND, not determined.

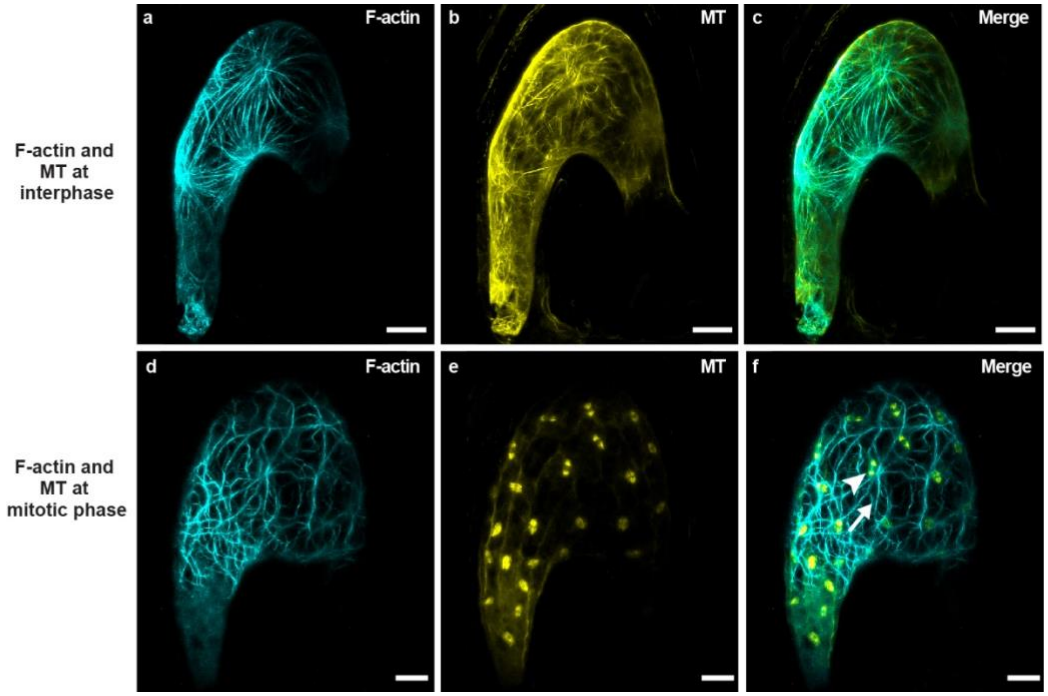


**Extended Data Figure 5: Disorganized reticulate F-actin at the moment when the MTs form spindles during mitotic nuclear division. a-f**, Time-lapse Z-projected confocal images of coenocytic endosperm F-actin (**a, d,** cyan, *proFWA::Lifeact:Venus*), MT (**b, e,** yellow, *proFWA::TagRFP::TUA5*), and both merged (**c, f**) before and/or after division (**a-c**) and at the onset of mitotic nuclear division (**d-f**). Scale bar, 20 µm. Arrowhead indicates MT spindle formation during mitotic nuclear division and arrow indicates disorganized reticulate F-actin in **f**.

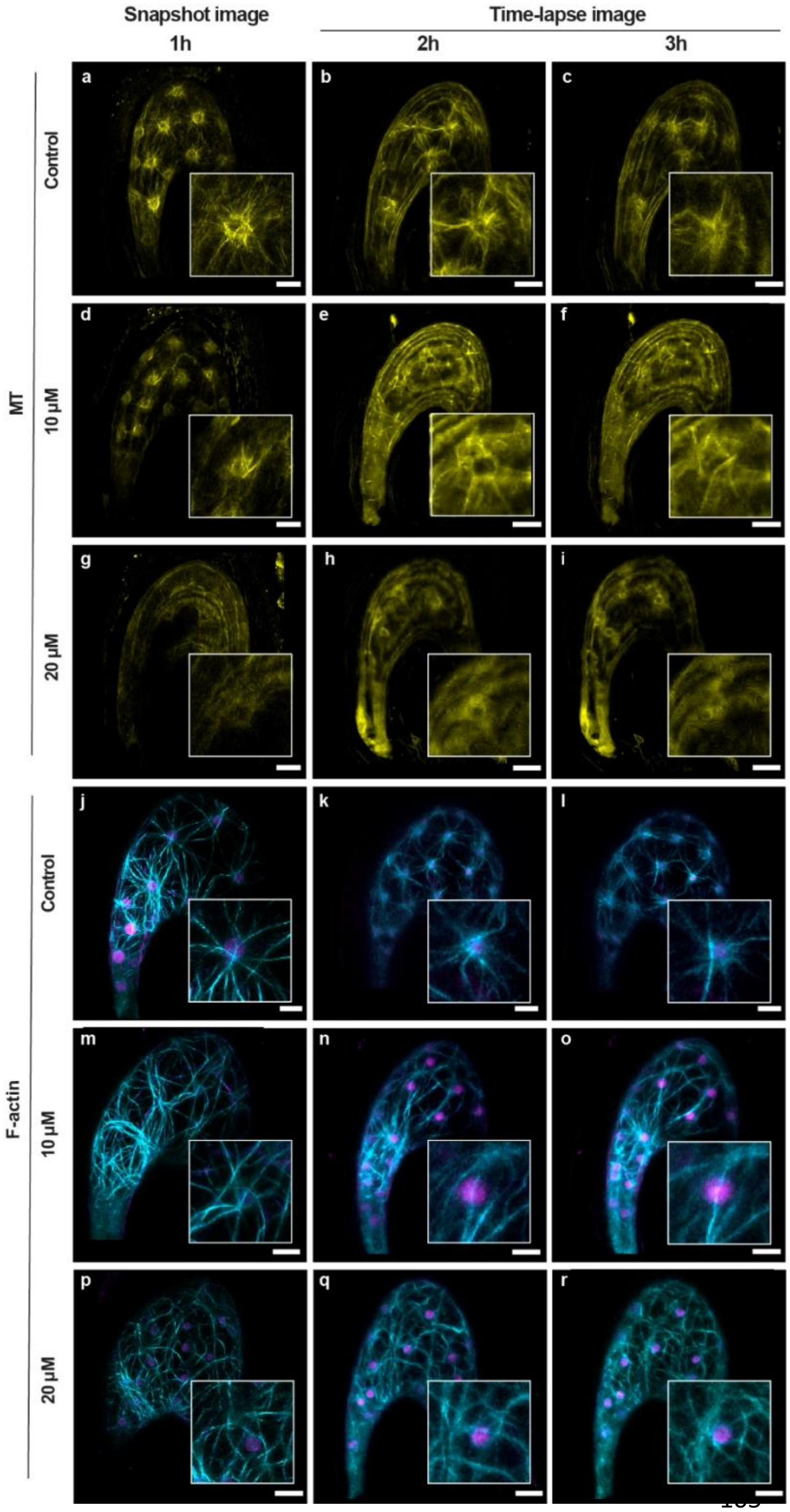


**Extended Data Figure 6: Dose-dependent effect of oryzalin to inhibit MT and F-actin aster structures. a-r,** Snapshot and time-lapse Z-projected confocal images of coenocytic endosperm MT (yellow, *proFWA::TagRFP::TUA5*), F-actin (cyan, *proFWA::Lifeact:Venus*), and nuclei (magenta, *proFWA::H2B:mRuby2*). MT control (**a-c**), MT 10 μM oryzalin (**d-f**), MT 20 μM oryzalin (**g-i**), F-actin control (**j-l**), F-actin 10 μM oryzalin (**m-o**), F-actin 20 μM oryzalin (**p-q**). Enlarged inserts at the right bottom of the images represent F-actin and MTs around the nucleus in **a-r**. Scale bar 20 μm.

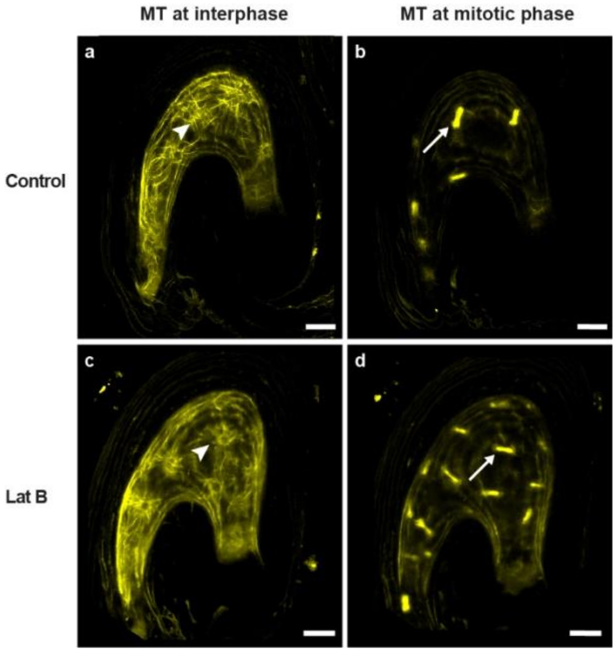


**Extended Data Figure 7: F-actin is not required for MT function during coenocytic endosperm development. a-d,** Time-lapse Z-projected confocal images of coenocytic endosperm MT (yellow, *proFWA::TagRFP::TUA5*) control (**a, b**), and 5 μM Lat B treatment (**c, d**). Lat B treatment did not affect MT organization (**c**) and nuclear division (**d**) during coenocytic endosperm development. Arrowheads indicate MT aster radiating from each nucleus and arrows indicate MT spindles formed during mitotic nuclear division. Scale bar, 20 μm.

**Video 1**: Combined time-lapse live-cell video showing 1st to 10th nuclear divisions in the *Arabidopsis* coenocytic endosperm. Endosperm nuclei marked by *proFWA::H2B:mRuby2* (magenta) (related to Figure 1). 1st seed (1st to 4th divisions, 20 min interval, 18 h in total); 2nd seed (4th to 6th divisions, 15 min interval, 29 h in total); 3rd seed (6th to 7th nuclear divisions, 20 min interval 29 h total); 4th seed (8th to 10th nuclear divisions, 20 min interval 44 h total). Note that cable structures are also visible in the 2nd and 4th seeds due to the simultaneous excitation of Lifeact-Venus *(proFWA::Lifeact-Venus)* by 514 nm laser. Scale bar = 20 µm.

**Video 2:** 3D structure of endosperm showing F-actin and endosperm nuclei visualized by *proFWA::Lifeact:Venus* (cyan) and *proFWA::H2B:mRuby2* (magenta), respectively in *Arabidopsis* coenocytic endosperm (related to Figure 2).

**Video 3:** Combined time-lapse live-cell video of F-actin dynamics and endosperm nuclei visualized by *proFWA::Lifeact:Venus* (cyan) and *proFWA::H2B:mRuby2* (magenta), respectively in *Arabidopsis* coenocytic endosperm (related to Figure 2). 1st seed (early stage of coenocytic endosperm, 15 min interval, 16 h in total, 2nd seed (late stage of coenocytic endosperm, 20 min interval, 27 h in total). Scale bar= 20 µm.

**Video 4:** Time-lapse (20 min interval) live-cell video of F-actin dynamics and endosperm nuclei visualized by *proFWA::Lifeact:Venus* (cyan) and *proFWA::H2B:mRuby2* (magenta), respectively in *Arabidopsis* coenocytic endosperm, control and 5 µM Lat B (related to Figure 2). Scale bar = 20 µm.

**Video 5:** Time-lapse (20 min interval) live-cell video (24 h in total) of F-actin dynamics and endosperm nuclei visualized by *proFWA::Lifeact:Venus* (cyan) and *proFWA::H2B:mRuby2* (magenta), respectively in *DN-ACTIN* (related to Figure 3). Scale bar = 20 µm.

**Video 6:** Time-lapse (20 min interval) live-cell video (44 h in total) of F-actin dynamics and endosperm nuclei visualized by *proFWA::Lifeact:Venus* (cyan) and *proFWA::H2B:mRuby2* (magenta), respectively in *OX-ACTIN* (related to Figure 3). Scale bar = 20 µm.

**Video 7:** 3D structure of coenocytic endosperm showing MTs visualized by *proFWA::TagRFP::TUA5* (yellow) in *Arabidopsis* coenocytic endosperm (related to Figure 4)**.**

**Video 8:** Time-lapse (20 min interval) live-cell video (24 h in total) of MT dynamics visualized by *proFWA::TagRFP::TUA5* (yellow) in *Arabidopsis* coenocytic endosperm of control, 10 µM oryzalin (related to Figure 5) and 5 µM Lat B (related to Extended Data Figure 7). Scale bar = 20 µm.

**Video 9:** Time-lapse (20 min interval) live-cell video (21 h in total) of F-actin dynamics and endosperm nuclei visualized by *proFWA::Lifeact:Venus* (cyan) and *proFWA::H2B:mRuby2* (magenta), respectively in *Arabidopsis* coenocytic endosperm of control, and 10 µM oryzalin (related to Figure 5). Scale bar = 20 µm.

**Video 10:** Time-lapse (30 min interval) live-cell video of MTs visualized by *proFWA::TagRFP::TUA5* (yellow), F-actin visualized by *proFWA::Lifeact:Venus* (cyan) and endosperm nuclei visualized *proFWA::H2B:mRuby2* (magenta) in *Arabidopsis* coenocytic endosperm of MT control, MT after oryzalin washout, F-actin control and F-actin after oryzalin washout (related to Figure 5). Scale bar = 20 μm.
